# *In planta* bacterial multi-omics analysis illuminates regulatory principles underlying plant-pathogen interactions

**DOI:** 10.1101/822932

**Authors:** Tatsuya Nobori, Yiming Wang, Jingni Wu, Sara Christina Stolze, Yayoi Tsuda, Iris Finkemeier, Hirofumi Nakagami, Kenichi Tsuda

**Author notes:** These authors contributed equally to this work (Tatsuya Nobori, Yiming Wang). Salk Institute for Biological Studies, La Jolla, CA 92037, USA. College of Plant Protection, Nanjing Agriculture University, Nanjing 210095, Jiangsu, China (Yiming Wang). National Key Laboratory of Plant Molecular Genetics, Shanghai Institute of Plant Physiology and Ecology, Chinese Academy of Sciences, 200032 Shanghai, China (Jingni Wu). Plant Physiology, Institute of Plant Biology and Biotechnology, University of Muenster, 48149 Münster, Germany (Iris Finkemeier).

## Abstract

Understanding how gene expression is regulated in plant pathogens is crucial for pest control and thus global food security. An integrated understanding of bacterial gene regulation in the host is dependent on multi-omic datasets, but these are largely lacking. Here, we simultaneously characterized the transcriptome and proteome of a foliar bacterial pathogen, *Pseudomonas syringae*, in *Arabidopsis thaliana* and identified a number of bacterial processes influenced by plant immunity at the mRNA and the protein level. We found instances of both concordant and discordant regulation of bacterial mRNAs and proteins. Notably, the tip component of bacterial type III secretion system was selectively suppressed by the plant salicylic acid pathway at the protein level, suggesting protein-level targeting of the bacterial virulence system by plant immunity. Furthermore, gene co-expression analysis illuminated previously unknown gene regulatory modules underlying bacterial virulence and their regulatory hierarchy. Collectively, the integrated *in planta* bacterial omics approach provides molecular insights into multiple layers of bacterial gene regulation that contribute to bacterial growth *in planta* and elucidate the role of plant immunity in controlling pathogens.

## Introduction

The growth of bacterial pathogens in plants and disease development are determined by genetically encoded bacterial virulence system and plant immune system (*1*). Despite the wealth of knowledge available concerning these systems in isolation, interactions between the two–especially how plant immunity affects bacterial function–are poorly understood (*2*). Previously, it was shown that *in planta* transcriptomics of a bacterial pathogen can be used to identify bacterial mRNAs whose expression is influenced by plant immune activation (*3*). Although transcriptome analysis is a useful and widely used approach to elucidate cellular function, it has been well established that mRNA expression does not always reflect protein expression, and thus it becomes clear that a better understanding of cellular behavior requires direct interrogation of protein expression (*4*, *5*). A previous study showed that proteome analysis of leaf commensal bacteria can reveal metabolic changes in bacteria residing on the leaf surface (*6*). However, the capacity of proteomics to describe plant-associated bacteria remains limited (*7*). For instance, analyzing bacterial responses in the intercellular space (apoplast) of leaves, which is an important niche for various commensal and pathogenic bacteria, poses a major challenge because the large preponderance of plant material relative to bacterial material confounds analysis. To date, there is no proteome study of bacterial pathogens in the leaf apoplast, and thus, we lack comprehensive knowledge of proteins that are affected by the host plants. Moreover, due to the lack of comparative analyses between different modalities of bacterial responses *in planta*, little is known about the flow of bacterial genetic information (mRNA and protein expression) that is important for virulence and how this is affected by plant immunity.

Here, we simultaneously profiled bacterial transcriptomes and proteomes *in planta* and identified bacterial processes influenced by plant immunity at the mRNA and protein levels at early and late stages of infection. Comparative analysis of transcriptomes and proteomes revealed that changes in bacterial mRNA and protein expression are correlated in general. However, proteins involved in the tip component of the bacterial type III secretion system were selectively suppressed by plant immunity at the protein level, implying the direct targeting of an essential virulence component by plant immunity. Furthermore, gene regulatory network analysis of bacteria showed previously unknown gene regulatory modules that mediate bacterial virulence *in planta*. Together, this study reveals the multi-layered regulatory mechanisms that underlie interactions between plants and bacterial pathogens.

## Results

### *In planta* transcriptome and proteome profiling of *P. syringae*

We profiled the transcriptome and proteome of the bacterial pathogen *Pseudomonas syringae* pv. *tomato* DC3000 (*Pto*) in *Arabidopsis thaliana* leaves using RNA-seq and liquid chromatography-mass spectrometry (LC-MS), respectively. Bacterial information was enriched by isolating bacterial cells from infected plant leaves using a previously established method (Fig. 1A) (*3*). Briefly, infected leaves were crushed and incubated in a buffer that stops bacterial metabolism and protects bacterial RNA from degradation. After bacterial cells and plant cells were separated by centrifugation, RNA and protein were extracted from isolated bacteria and subjected to RNA-seq and LC-MS analysis, respectively (Fig. 1A). Both the transcriptomes and proteomes of *Pto* were profiled under 15 conditions, including two *in vitro* conditions and 13 *in planta* conditions using combinations of five *A. thaliana* genotypes, two *Pto* strains, and two time points (Fig. 1B). In the proteome analysis, we could detect up to approximately 2,000 proteins *in planta* (Fig. 1C). The numbers of mRNA/proteins detected in different subcellular locations were proportional to the number of *Pto* genome copies regardless of conditions (Fig. 1C), suggesting that no obvious bias was introduced into mRNA/protein detection during bacterial enrichment processes. *Pto* showed distinct responses under different conditions at both transcriptome (Fig. 1D and 1E) and proteome (Fig. 1F and 1G) levels. *In planta* transcriptome and proteome patterns were more distinct from *in vitro* data at 6 h post infection (hpi) compared with 48 hpi (Fig. 1 D-G), implying that dynamic transcriptional reprogramming at an early stage of infection is crucial for *Pto* to adapt to the plant apoplastic environment and become virulent. This notion is supported by our previous observation that bacterial transcriptome patterns at this early stage of infection can predict bacterial growth in plants at a later stage (*8*).

**Fig. 1:**
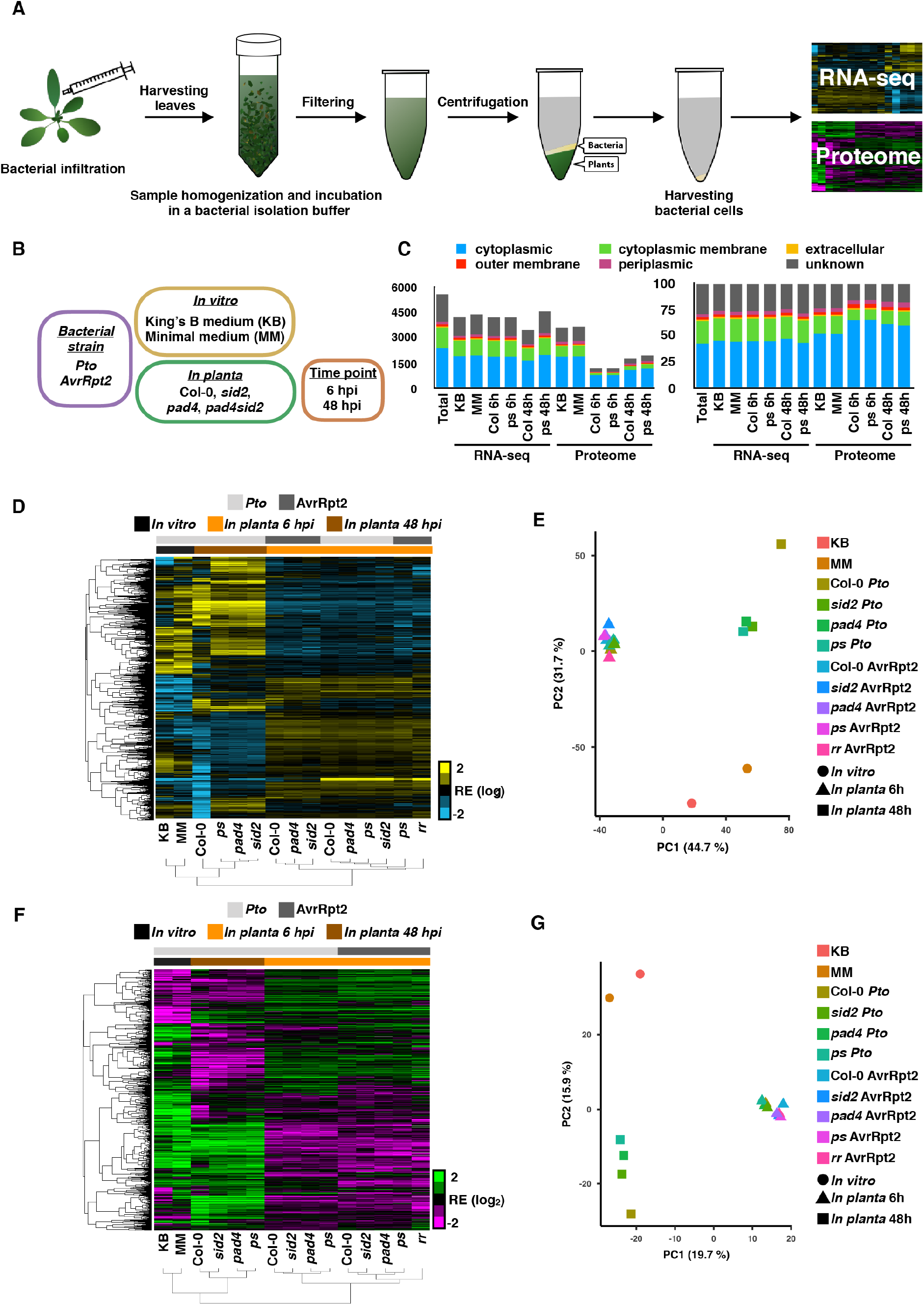
*In planta* transcriptomics and proteomics of *Pto*. (A) Schematic workflow of *in planta* bacterial transcriptome and proteome analysis. (B) Bacterial strains, plant genotypes, and conditions used in this study. (C) Number (left) and proportion (right) of mRNAs/proteins detected in the RNA-seq/proteome analysis together with protein localization information. Total: all annotated genes of *Pto*. *ps*: *pad4 sid2* (D) Hierarchical clustering of relative expression (RE) of commonly detected *Pto* and *Pto* AvrRpt2 genes (4,868 genes) in RNA-seq analysis. See Data S1 for the gene expression data. (E) Principle component analysis of the *Pto* and *Pto* AvrRpt2 genes commonly detected in RNA-seq analysis. (F) Hierarchical clustering of the relative expression of *Pto* and *Pto* AvrRpt2 proteins commonly detected in LC-MS analysis (937 proteins). See Data S2 for the expression of proteins detected in at least one condition (2,018 proteins). (G) Principle component analysis of *Pto* and *Pto* AvrRpt2 proteins commonly detected in LC-MS analysis.

### Dynamic regulation of bacterial function across different conditions

Many biological processes of *Pto* were differentially regulated in distinct conditions (Fig. S1). To gain further insights into these biological functions, we grouped *Pto* genes into gene ontology (GO) terms and calculated standardized GO expression scores; GO terms expressed in highly condition-dependent manners were then selected (Fig. 2A and 2B; see Materials and Methods). The GO term “pathogenesis” was one of the most dynamically regulated processes at both the transcriptome and proteome levels (Fig. 2A and 2B). These mRNAs and proteins were strongly induced *in planta* at 6 hpi; their expression remained high at 48 hpi, and a clear host genotype effect was observed at this time point; i.e., expression was higher in the mutants of the salicylic acid (SA) pathway (*sid2*, *pad4*, *pad4 side2*) compared with the wild type Col-0 (Fig. 2C and 2D), suggesting that SA-mediated immunity suppresses pathogenesis-related factors at the transcript level at 48 hpi. *Pto* AvrRpt2 induces effector-triggered immunity (ETI) upon recognition of the effector AvrRpt2 by the receptor RPS2 (*9*, *10*), and SA pathways are important components of this ETI (*11*). We found that successful activation of ETI strongly induces mRNAs/proteins related to “catalase activity” at 6 hpi (Fig. S2B), which probably reflects bacterial responses to oxidative burst, a hallmark of ETI responses. Interestingly these mRNAs and proteins were even more highly expressed in virulent *Pto* at 48 hpi (Fig. S2B), implying that *Pto* experiences oxidative stress at later infection stages. Taken together, our multi-omic *Pto* dataset uncovered dynamic regulation of various biological processes across different conditions at the mRNA and protein levels.

**Fig. 2:**
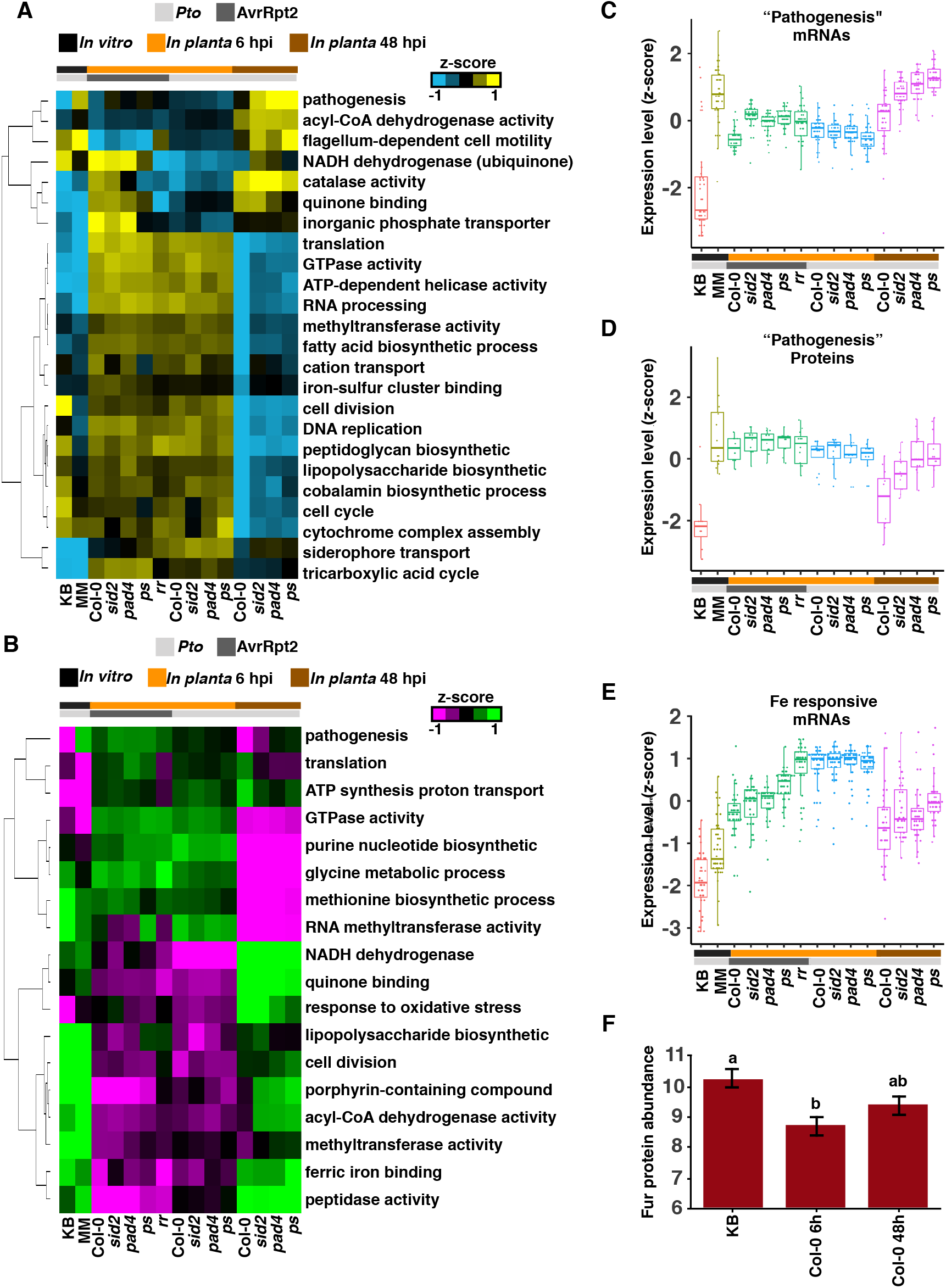
Dynamic regulation of bacterial function across different conditions. (A and B) Hierarchical clustering of selected gene ontology (GO) terms in transcriptome (A) and proteome (B) data. Transcriptome and proteome data were standardized using z-scores (log_2_) and mean z-scores of mRNAs/proteins involved in individual GO terms were shown. For the full GO expression data, see Data S7 and S8. (C and D) Box plots show expression (z-score) of mRNAs (C) and proteins (D) related to “pathogenesis”. (E) Box plots show expression (z-score) of mRNAs previously described as “iron-responsive” (*13*). (A-E) Light and dark gray sidebars represent *Pto* and *Pto* AvrRpt2, respectively. Black, orange, and brown sidebars represent *in vitro* (KB), *in planta* (Col-0) 6 hpi, and *in planta* 48 hpi, respectively. (F) Expression of the Fur protein in *Pto* based on the proteome data (normalized iBAQ value (log_2_)). Different letters indicate statistically significant differences (adjusted *p*-value < 0.01; Benjamini–Hochberg method).

Previously, we showed that bacterial genes related to the iron acquisition pathway (iron starvation genes) are highly induced in susceptible plants and strongly suppressed by the activation of pattern-triggered immunity (PTI) and/or ETI at 6 hpi at the mRNA level (*3*) (Fig. 2E). Expression of these genes was lower at 48 hpi compared with 6 hpi (Fig. 2E), but still higher than under *in vitro* conditions. Expression of iron starvation genes is known to be regulated by the master regulator protein Fur, which typically functions as a transcriptional repressor when bound by Fe(II) (*12*). Interestingly, the accumulation of Fur protein was negatively correlated with the expression of iron starvation genes in three distinct conditions (*in vitro*, *in planta* 6 hpi, and *in planta* 48 hpi) (Fig. 2E and 2F). This implies a previously unknown mechanism of bacterial iron acquisition, by which accumulation of the Fur protein might also contribute to regulation of iron-starvation genes.

### Comparative analysis of bacterial transcriptomes and proteomes

To compare global expression patterns of genes and proteins, the transcriptome and proteome data of all 15 conditions were standardized and combined, and hierarchical clustering was performed (Fig. S3A). Strikingly, transcriptome and proteome data were clustered together in three major conditions (*in vitro* and *in planta* 6/48 hpi) (Fig. S3A), indicating that the global patterns of bacterial gene expression and protein expression are similar both *in vitro* and *in planta*. Since many of the samples for transcriptome and proteome data were prepared independently, the overall agreement between transcriptome and proteome data indicates the high accuracy of both sets of omics data.

We compared RNA-seq and proteome data in each condition. In King’s B medium (KB) and minimal medium (MM) conditions, transcriptomes and proteomes were moderately correlated (R^2^ = 0.52 and 0.43, respectively) (Fig. S3B), which is consistent with previous studies in *Escherichia coli* (R^2^ = 0.42 − 0.57) (*14*–*16*). A similar level of correlation was observed *in planta* with slightly higher correlation at 6 hpi (R^2^ = 0.51 − 0.55) than at 48 hpi (R^2^ = 0.39 − 0.47) (Fig. S3B). Thus, *Pto* mRNA and protein expression are moderately correlated both *in vitro* and *in planta*.

We further compared the fold changes in the RNA-seq and proteome data between *Pto in vitro* (KB) and each of the other conditions (Fig. 3A). In all conditions, expression changes in transcriptome and proteome were moderately correlated (R^2^ = 0.52 − 0.63), suggesting that protein expression changes closely mirror changes in bacterial mRNA levels during infection to both resistant and susceptible plants. Of 1,068 mRNAs/proteins detected in both RNA-seq and proteome analyses, 111 mRNAs/proteins (10.4%) were significantly induced at both transcriptome and proteome levels *in planta* at 6 hpi compared with in KB (Fig. 3B). GO analysis showed that “pathogenesis-related process” was enriched among these mRNAs/proteins (Fig. 3B), indicating the transcription-driven activation of pathogenesis programs upon plant infection. On the other hand, there were cases where expression of mRNAs and proteins were discordant. Interestingly, more proteins were down-regulated (168 proteins) than up-regulated (39 proteins) in a protein-specific manner (Fig. 3B). This may be explained by a prominent role of protein degradation or translation inhibition in bacteria *in planta* (Fig. 3B). GO analysis showed that “cell wall biogenesis”-related proteins were suppressed only at the protein level (Fig. 3B). In contrast, more mRNAs were up-regulated (207 proteins) than down-regulated (49 proteins) in an mRNA-specific manner (Fig. 3B). This implies that upregulation of specific mRNAs is a key response of *Pto* at an early stage of infection, and that the induction of mRNAs is not yet reflected in protein abundance at this point. We also compared the fold changes in the transcriptome and proteome profiles between *pad4 sid2* and Col-0 at 48 hpi. GO enrichment analysis showed that bacterial processes related to “chemotaxis” were highly expressed in *pad4 sid2* at the protein level, but not the mRNA level (Fig. S3C). This suggests that chemotaxis-related processes are suppressed by plant SA-mediated immunity at the protein level, and this may be important for bacterial growth inhibition. Collectively, genome-wide comparisons between mRNA and protein expression illuminate the multifaceted control of bacterial gene expression *in planta*.

**Fig. 3:**
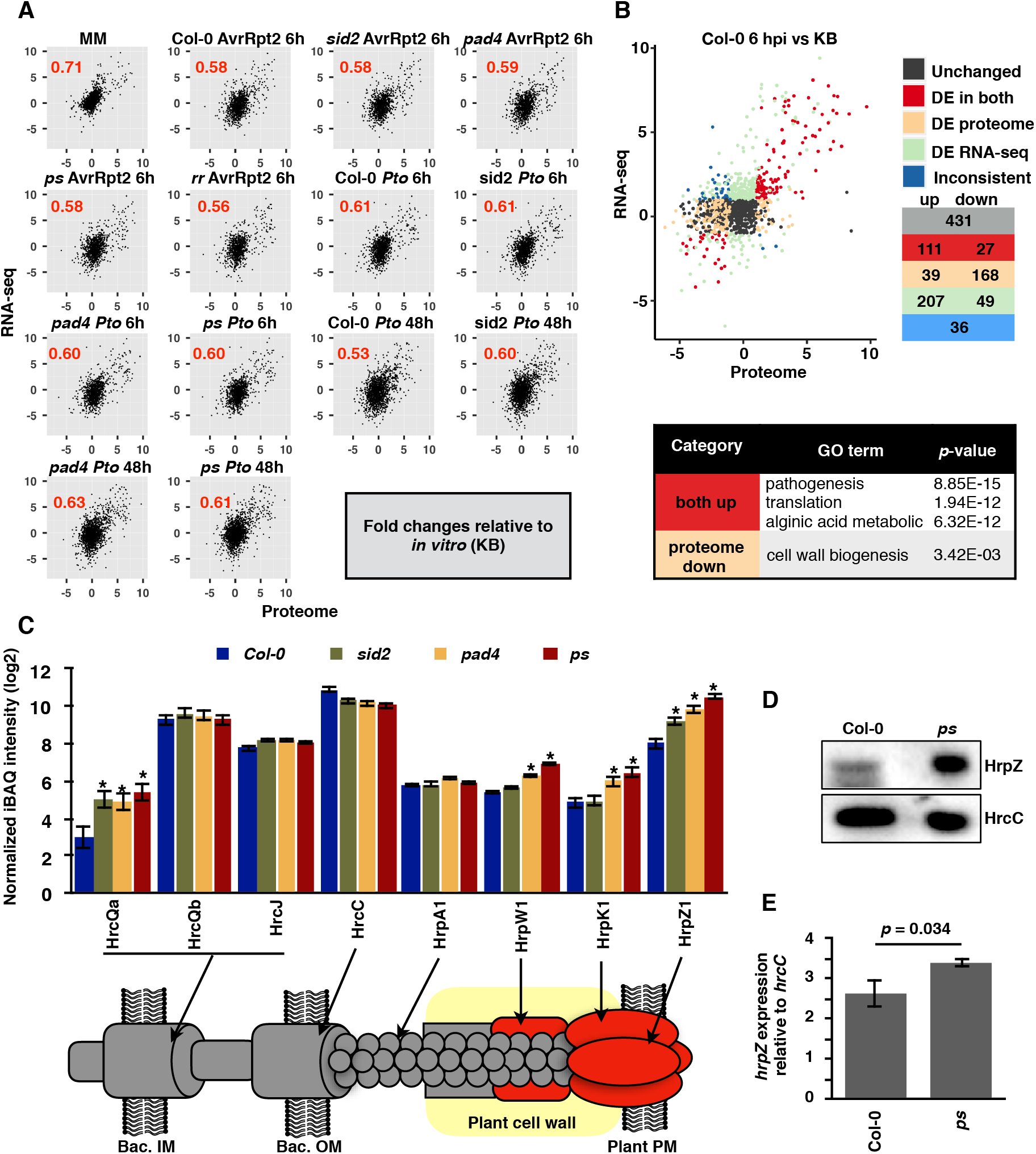
Conserved and divergent regulation of mRNA and protein expression in *Pto.* (A) Comparisons between transcriptome and proteome data in each condition. Fold changes relative to *in vitro* (KB) conditions were subjected to analysis. Pearson’s correlation coefficients are shown. (B) mRNA/protein expression fold changes between *in planta* (Col-0) at 6 hpi and *in vitro* (KB) were compared. mRNAs/proteins differentially expressed (DE; FDR < 0.01; |log_2_FC| | 2) in both, either, and neither (“Unchanged”) transcriptome and proteome studies were grouped and colored. mRNAs/proteins differentially expressed in the opposite direction are colored in blue (“Inconsistent”). The numbers of mRNAs/proteins are shown for each category. List of gene ontology (GO) terms enriched in the group of proteins that are significantly induced *in planta* at both mRNA and protein levels or proteins that are significantly suppressed *in planta* only at the protein level. For the full gene list and GO list, see Data S10 and S11. (A and B) mRNAs/proteins detected in both the transcriptome and proteome in each comparison were used for this analysis. (C) Expression of T3SS-related proteins based on the proteome data (normalized iBAQ value). Approximate localization of each protein is shown in the diagram. (D) HrpZ and HrcC proteins detected by immunoblotting using specific antibodies. Protein loading amount was adjusted relative to the expression of the HrcC protein. Similar results were obtained in three independent experiments. (E) RT-qPCR analysis of *hrpZ* and *hrcC* relative to 16S. Plants were infiltrated with *Pto* at OD_600_ = 0.05 and harvested at 48 hpi.

### Component-specific targeting of the type III secretion system by plant SA pathways

GO enrichment analysis showed that the plant SA pathway suppresses a significant number of bacterial proteins related to pathogenesis including proteins comprising the type III secretion system (T3SS) (Fig. S1D). The T3SS is an essential component by which *Pto* translocates effectors into plant cells to subvert plant immunity and become virulent (*17*). We found that the impact of the SA pathways was apparent almost exclusively in proteins comprising the tip of the T3SS, namely HrpZ, HrpK, and HrpW (Fig. 3C). This implies that the SA pathways selectively target the tip of bacterial T3SS. To confirm this observation, we performed immunoblotting using protein directly extracted from infected leaves without physical bacterial isolation. When protein loading was normalized by HrcC expression, HrpZ accumulated more highly in *pad4 sid2* plants than in Col-0 plants, indicating differential effects of the SA pathway on different components of the type III secretion system (Fig. 3D and Fig. S4A). This also suggests that the bacterial isolation process did not introduce artefacts in the proteome data. To test if differential expression of HrpZ is due to different bacterial populations in plants, we compared HrpZ protein abundance between Col-0 at 48 hpi and *pad4 sid2* at 24 hpi, time points at which bacterial population densities were comparable (Fig. S2C). Also in this comparison, HrpZ protein accumulated to higher levels (1.6-fold) in *pad4 sid2* than in Col-0 (Fig. S4B), suggesting that bacterial population does not solely explain differences in HrpZ protein expression and thus that SA-mediated immunity might directly target this protein. Furthermore, the difference in relative protein accumulation (*hrpZ/hrcC*) between Col-0 and *pad4 sid2* at 48 hpi (5.9-fold) could not be solely explained by differences in mRNA expression (1.7-fold) (Fig. S4B and Fig. 3E). Taken together, the results suggest that SA-mediated immunity selectively affects expression of the tip component of the T3SS at the protein level.

### Gene co-expression analysis predicts bacterial gene regulatory logic

Despite the distinct regulation of certain specific mRNAs and proteins, the overall moderate correlation between transcriptome and proteome patterns of *Pto* suggests that mRNA expression can be a good indicator of bacterial functional expression *in planta*. Also, transcriptome-based analysis would be aided by additional transcriptome data available from a previous study (*8*) and by the fact that a greater number of *Pto* mRNAs were detected compared to proteins (Fig. 1C). Therefore, we reasoned that investigating the regulatory network governing bacterial mRNA expression would help deepen our understanding of bacterial functional regulation. To deconvolute the gene regulatory network of *Pto*, we used 125 transcriptome datasets of *Pto* profiled in 38 conditions (generated in a previous study (*3*) and this study). A correlation matrix of 4,765 genes revealed highly correlated gene clusters (Fig. S5A), some of which were enriched with known functions (Fig. 4A). Then, we built a gene co-expression network based on the correlation scores and annotated genes with known functions; this allowed us to conclude that genes sharing the same functions tend to be co-expressed (Fig. 4B). For instance, genes related to pathogenesis (mostly T3SS and effector genes), flagellum, and iron starvation responses were found in separate and highly co-expressed gene clusters (Fig. 4A and 4B). Intriguingly, genes involved in coronatine biosynthesis and alginate biosynthesis were clustered very closely together, suggesting that these processes might share the same regulatory mechanism (Fig. S5B). On the other hand, genes related to coronatine biosynthesis and the T3SS were only mildly correlated with each other (Fig. S5B), although it has been shown that the expression of *corR*, the master regulator of coronatine biosynthesis genes, is dependent on HrpL, the master regulator of the T3SS (*18*). This suggests that there might be additional regulators that govern the expression patterns of genes related to coronatine biosynthesis and the T3SS. We also found that some genes annotated as effectors were not co-expressed with the majority of effectors (Fig. S5B), suggesting that they function in different contexts or that they do not function as effectors. Strong anti-correlation was observed between “siderophore transport” genes, which are iron-repressive, and “ferric iron binding” genes, which are involved in iron-inducible bacterioferritin (Fig. S5C), indicating that this analysis could capture known expression patterns.

**Fig. 4:**
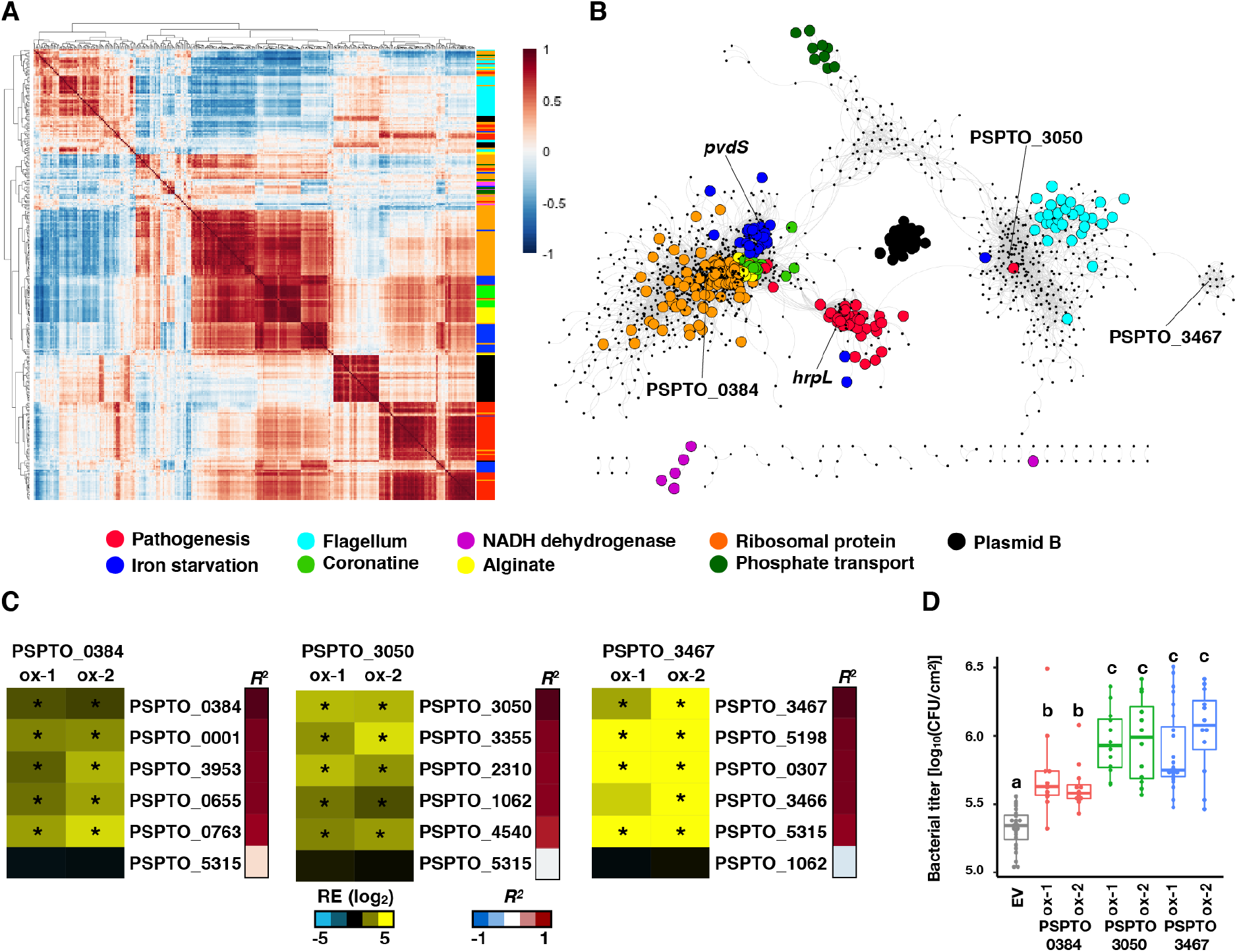
mRNA co-expression analysis of *Pto* reveals functional mRNA modules. (A) Correlation matrix of selected *Pto* mRNAs (330 mRNAs) with functional annotations shown in the side bar. For the full dataset, see Fig. S5A and Data S13. (B) mRNA co-expression network of *Pto*. Each node represents an mRNA. Highly correlated (*R*^2^ > 0.8) mRNAs are connected with edges. Functions are annotated with colors. (C) Two independent *Pto* strains overexpressing putative transcriptional regulators (TRs) were generated. RT-qPCR analysis of *in vitro* mRNA expression fold changes between *Pto* overexpressing putative TRs and wild type *Pto*. Asterisks indicate significant difference (|log_2_FC| > 1, *p*-value < 0.05). Correlation coefficients between each mRNA and the TRs are shown in the sidebars on the right. (D) Growth of *Pto* overexpressing putative TRs or an empty vector (EV) (infiltrated at OD_600_ = 0.001) in Col-0 at 48 hpi. n = 12 biological replicates from three independent experiments. Different letters indicate statistically significant differences (adjusted *p*-value < 0.01; Benjamini–Hochberg method).

We anticipated that groups of highly co-expressed genes contain transcriptional regulators (TRs) and their targets. Indeed, *hrpL* and the iron starvation sigma factor *pvdS* were co-expressed with their targets, T3SS genes and iron starvation genes, respectively (Fig. 4B). To test whether gene co-expression data can predict gene regulatory hierarchy, we selected three putative TRs, PSPTO_0384, PSPTO_3050, and PSPTO_3467, whose functions have not been characterized, and generated *Pto* strains that overexpress each of the TRs. We then analyzed the expression of predicted target genes which were highly co-expressed with individual TRs. Remarkably, for the three TRs, all or most of predicted target genes were highly expressed in the TR overexpression lines *in vitro* (Fig. 4C), supporting the predicted regulatory hierarchy. This was further confirmed *in planta* at 6 hpi for PSPTO_3050, but overexpression of PSPTO_0384 or PSPTO_4908 induced only a small number of the predicted targets (Fig. S6A). This is probably explained by the fact that genes highly co-expressed with these two TRs are already strongly induced in wild type *Pto* at 6 hpi (Fig. S6B), and thus, overexpression of these TRs did not lead to further induction of the predicted target genes. Notably, all the TR-overexpressing *Pto* grew significantly better than wild type *Pto* in Col-0 plants (Fig. 4D and Fig. S6C), suggesting that the three TRs and some of their regulons play positive roles in *Pto* growth in plants. Collectively, *in planta* transcriptome data of *Pto* isolated from hosts with diverse genotypes at different time points enables gene co-expression analysis that can be used to identify previously unknown bacterial gene clusters that contribute to bacterial growth *in planta* as well as to reveal the regulatory logic in the gene clusters.

## Discussion

In this study, we analyzed the transcriptome and proteome of the bacterial pathogen *Pto* both *in vitro* and *in planta* under various conditions. This is, to our knowledge, the first study integrating transcriptomes and proteomes of bacteria in a plant host. We found that bacterial mRNA and protein expression were moderately correlated in liquid media and in resistant and susceptible plants (Fig. 3A). Our data indicate that population-level changes in bacterial transcriptomes can serve as a reliable predictor of the proteome changes elicited by plant colonization. Previous studies using plants, yeasts, and mammals showed varying degrees of correlation between transcriptomes and proteomes, but in most cases, the correlation was considerably lower than for the bacterial pathogen *Pto* shown in this study (*19*–*23*). This suggests that, in *Pto* at least, at the population level, mRNAs are faithfully translated into proteins. On the other hand, a previous study showed that, at the single cell level, mRNA and protein levels are completely uncorrelated in another Gram-negative bacterium, *E. coli*, grown in a microfluidic device (*24*). Single cell analysis will be required to determine the degree of correlation between the transcriptome and proteome in individual cells of *Pto in planta*.

Multimodal measurements of bacterial responses during infection provide a systematic view of bacterial gene regulation that cannot be captured by analyses of any single modality alone. By analyzing cases in which mRNA and protein expression do not correlate with each other, we found that the SA pathway of *A. thaliana* selectively suppresses expression of the tip component of the T3SS at the protein level (Fig. 3C–3E). These proteins, HrpZ1, HrpW1, and HrpK, were shown to function redundantly in type III effector (T3E) translocation (*25*), and lack of *hrpK* alone compromised growth of *Pto* in *A. thaliana* (*26*). Thus, targeting these proteins is sufficient to dampen the virulence of *Pto*. Our results imply that SA-mediated plant immunity might promote the degradation of these proteins to suppress T3E translocation into plant cells and thus to inhibit pathogen growth. It is tempting to speculate that plant immunity selectively and directly targets the tip component of the T3SS at the plant cell wall. As the T3SS must cross the cell wall to successfully translocate T3Es into plant cells, it therefore represents an ideal target for plants to counter bacterial virulence. The mechanism by which plants target the tip component of the T3SS awaits further investigation. mRNA co-expression analysis identified groups of highly co-expressed bacterial mRNAs with known and unknown functions. Using mRNA co-expression data, we could predict relationships between three putative TRs and mRNAs whose expression is affected by these TRs (Fig. 4C). The TRs and their potential target mRNAs are induced *in planta* at an early and/or a late time point of infection (Fig. S6B) and overexpression of the TRs led to enhanced bacterial growth *in planta* (Fig. 4D). Therefore, our approach has the capability to identify previously unknown gene modules that are important for virulence in plants. mRNAs co-expressed with the TR PSPTO_0384 are suppressed by plants that have engaged PTI (flg22 pre-treatment) (Fig. S6B), which is similar to known virulence related genes such as the T3SS and effectors (*3*). Thus, the induction of these mRNAs might be important for virulence and this might explain the enhanced growth of PSPTO_0384-ox strains *in planta* (Fig. 4D). Interestingly, mRNAs co-expressed with either of the other two TRs PSPTO_0350 or PSPTO_3467 were induced by PTI activation, while overexpression of these TRs led to enhanced bacterial growth (Fig. 4D). It is possible that these genes are involved in stress adaptation and contribute to bacterial growth in plants. Previously, we observed that PTI activation induces a number of mRNAs in *Pto* (800 mRNAs) whose functions are not well understood (*3*). Investigating such genes might help us identify previously uncharacterized genes related to bacterial stress adaptation and/or virulence in plants. Taken together, *in planta* bacterial multi-omics represents a new strategy for studying the molecular mechanisms underlying bacterial virulence and plant immunity.

## Acknowledgements

We thank Sheng Yang He for providing the HrpZ antibody, Hsiou-Chen Huang for providing the HrcC antibody, Anne Harzen and Katharina Kramer for support in sample preparation and mass spectrometry analysis, Alan Collmer for the pCPP5040 plasmid, the Max-Planck Genome Centre for sequencing support, and Neysan Donnelly for critical comments on the manuscript. This work was supported by Huazhong Agricultural University, the Max Planck Society, a German Research Foundation grant (SPP2125) (to K.T.), a predoctoral fellowship from the Nakajima Foundation (to T.N.), and a postdoctoral fellowship from the Alexander von Humboldt Foundation (to Y.W.).

## Author contributions

T.N. and K.T. designed the research. T.N., Y.W., J.W., S.C.S., Y.T., I.F., and H.N. performed experiments. T.N. and K.T. wrote the paper with input from all authors.

## Data availability

The RNA sequencing data used in this study are deposited in the National Center for Biotechnology Information Gene Expression Omnibus database (accession no. GSE138901). The mass spectrometry proteomics data are available at the ProteomeXchange Consortium via the PRIDE partner repository with the data set identifier PXD015839.

## Conflict of interest

The authors declare no conflict of interest.

## Materials and Methods

### Plant materials and growth conditions

The *Arabidopsis thaliana* accession Col-0 was the background of all *A. thaliana* mutants used in this study. The *A. thaliana* mutants *rpm1-3 rps2-101C* (*27*), *pad4-1* (*28*), *sid2-2* (*29*), and *pad4 sid2* (*11*) were described previously. Plants were grown in a chamber at 22°C with a 10-h light period and 60% relative humidity for 24 days and then in another chamber at 22°C with a 12-h light period and 60% relative humidity. For all experiments, 31- to 33-day-old plants were used.

### Bacterial strains

*Pto* DC3000 carrying an empty vector (pLAFR) or *avrRpt2* (pLAFR) (*30*) were described previously. The *Pto* overexpression strains were generated as previously described (*31*). The coding sequences of *GFP*, *PSPTO_0384*, *PSPTO_3050*, and *PSPTO_3467* were amplified by PCR, cloned into the pENTR/D-TOPO vector, and then transferred into pCPP5040 (gentamicin-resistant) by the LR reaction. The *Pto* overexpression strains were generated by a triparental mating of the *Pto* wild type strain, *E. coli* carrying each construct, and an *E. coli* strain carrying pRK2013 (kanamycin-resistant). The transformed *Pto* strains were selected with 50 μg ml^−1^ rifampicin, 5 μg ml^−1^ gentamicin, and 50 μg ml^−1^ kanamycin.

### Accession numbers

The accession numbers for the genes discussed in this article are as follows: *AtPAD4* (AT3G52430), *AtSID2* (AT1G74710), *AtRPS2* (AT3G03600), *AtRPM1* (AT3G07040), *hrcC* (PSPTO_1389), and *hrpZ* (PSPTO_1382).

### Preparation of *in vitro* bacterial samples

Bacteria were grown in either King’s B medium or type III-inducible medium (*32*) (50 mM KH_2_PO_4_; 7.6 mM (NH_4_)_2_SO_4_; 1.7 mM NaCl; 1.7 mM MgCl_2_6H_2_O; 10 mM fructose) at 28°C until they reached OD_600_ = 0.65 (exponential phase). Upon harvesting the bacterial culture, 0.1 volumes of 5% phenol and 95% ethanol were added. The culture was then centrifuged to harvest the bacterial pellet, followed by total RNA and/or protein extraction.

### Bacterial infection of plant leaves and sampling

*Pto* stains were cultured in King’s B medium at 28°C at 200 rpm. Bacteria were harvested by centrifugation and resuspended in sterile water to OD_600_ of 0.5 (~2.5 × 10^8^ cfu/mL) and 0.005 (~2.5 × 10^6^ cfu/mL) for harvesting at 6 hpi and 48 hpi, respectively. In total, 80–100 *A. thaliana* leaves (four leaves per plant) were syringe-inoculated with bacterial suspensions using a needleless syringe. The infected leaves were harvested at 6 hpi or 48 hpi, immediately frozen in liquid nitrogen, and stored at −80°C. The bacterial growth assay was performed as described before (*33*).

### *In planta* bacterial transcriptomics

#### Sample preparation and RNA sequencing

*In planta* bacterial transcriptome analysis was conducted as described previously (*34*). Briefly, bacterial cells were isolated from plant leaves, followed by RNA extraction, DNase treatment, rRNA depletion, and cDNA library preparation. The cDNA libraries were sequenced using an Illumina HiSeq 3000 system with a 150-bp strand-specific single-end read, resulting in ~10 million reads per sample. The resulting reads were mapped onto the *Pto* DC3000 genome/CDS (*Pseudomonas* Genome Database) using Bowtie2 (*35*). Mapped reads were counted with the Python package HTSeq (*36*). The RNA-seq data used in this study are deposited in NCBI Gene Expression Omnibus database (accession no. GSE138901).

#### Data analysis

The statistical analysis of the RNA-seq data was performed in the R environment. Genes with average counts < 5 were excluded from the analysis. The count data were normalized and log-transformed by the function calcNormFactors [trimmed mean of M-values (TMM) normalization] in the package edgeR and the function voomWithQualityWeights in the package limma, respectively. To each gene, a linear model was fitted by using the function lmFit in the limma package with the following terms: Sgtr = GTgt + Rr + egtr, where S is the log_2_ count per million, GT is the host genotype:*Pto* strain interaction and the random factors, R is the biological replicate, and e is the residual. The eBayes function in the limma package was used for variance shrinkage during the calculation of the P values. The false discovery rate (FDR; the Storey’s q-values) was calculated using the qvalue function in the qvalue package (*37*). Genes with q-value <0.01 and log_2_ fold change > 2 were defined as differentially expressed genes. The prcomp function was used for principal component analysis. Hierarchical clustering was performed using the dist and hclust functions in the R environment or using Cluster3.0 software (*38*). Heatmaps were created with the heatmap3 or pheatmap function in the R environment or using TreeView (*39*). Enriched GO terms were identified using the BiNGO plugin for Cytoscape (*40*). Scatterplots and boxplots were generated using the R-package ggplot2. Correlation matrices were made by cor function and the correlation heatmap was drawn by pheatmap in the R environment. Gene correlation networks were created in Cytoscape with the yFiles Layout Algorithm.

#### Gene co-expression analysis

RNA-seq data obtained in a previous study (*8*) and the present study were combined (see Data S12 for the full dataset). Data were TMM-normalized and voom- (log_2_) transformed. Pair-wise Pearson’s correlation coefficients were calculated. Keywords related to particular functions were searched for on the gene annotation (available at www.pseudomonas.com) to stratify genes for the sake of visualization (Fig. 4A). Keywords used in this analysis were as follows: “type III”, “alginate”, “flagellar”, “flagellin”, “flagellum”, “NADH dehydrogenase”, “coronamic acid”, “coronafacic acid”, “ribosomal protein”, “phosphate transporter”, and “PSPTO_B”. Iron starvation genes were selected based on a previous study (*13*).

### *In planta* bacterial proteomics

#### Bacterial isolation and protein extraction

Bacterial isolation was done as described above. The TriFast solution was mixed with 0.2 volume of chloroform and the organic (lower) phase was isolated by centrifugation. The aqueous (upper) phase was isolated for RNA extraction for simultaneous profiling of transcriptomes and proteomes. The organic phase was mixed with 4 volumes of MeOH 0.01M ammonium acetate and incubated at −20°C overnight to precipitate proteins. The precipitated proteins were washed twice with MeOH 0.01 M ammonium acetate and then washed once with 80% acetone. Proteins were stored in 80% acetone at −20°C.

#### Sample preparation and fractionation

Proteins were pelleted and re-dissolved in 8 M urea 100 mM Tris-HCl pH 8.5, and then protein mixtures were reduced with dithiothreitol, alkylated with chloroacetamide, and digested first with Lys-C for 3 h and subsequently with trypsin o/n. Samples were submitted to SDB-RPS fractionation using a protocol adapted from a previous report (*41*). In brief, stage tips were prepared with two layers of SDB-RPS membrane and activated with 100 μL acetonitrile, followed by equilibration with 100 μL equilibration buffer (30% (v/v) MeOH, 1% (v/v) trifluoroacetic acid (TFA)) and 100 μL 0.2% TFA. Next, peptides were immobilized on the membrane and washed with 100 μL 0.2% TFA. Peptides were then eluted into three consecutive fractions using SDB-RPS buffer 1 (100 mM NH_4_HCO_2_, 40% (v/v) ACN, 0.5% FA), SDB-RPS buffer 2 (150 mM NH_4_HCO_2_, 60% (v/v) acetonitrile, 0.5% formic acid) and finally SDB-RPS buffer 3 (5% Ammonia (v/v), 80% (v/v) acetonitrile). The collected fractions were evaporated to dryness to remove residual ammonia.

#### LC-MS/MS data acquisition

Dried peptides were re-dissolved in 2% ACN, 0.1% TFA for analysis and adjusted to a final concentration of 0.1 μg/μl. Samples were analyzed using an EASY-nLC 1200 (Thermo Fisher) coupled to a Q Exactive Plus mass spectrometer (Thermo Fisher) or using an EASY-nLC 1000 (Thermo Fisher) coupled to a Q Exactive mass spectrometer (Thermo Fisher), respectively. Peptides were separated on 16-cm frit-less silica emitters (New Objective, 0.75 μm inner diameter), packed in-house with reversed-phase ReproSil-Pur C18 AQ 1.9 μm resin (Dr. Maisch). Peptides (0.5 μg) were loaded on the column and eluted for 115 min using a segmented linear gradient of 5% to 95% solvent B (80% ACN, 0.1%FA) (0–5 min: 5%, 5–65 min: 20%, 65–90 min: 35%, 90–100 min: 55%, 100–115 min: 95%) at a flow rate of 300 nL/min. Mass spectra were acquired in data-dependent acquisition mode with the TOP15 method. MS spectra were acquired in the Orbitrap analyzer with a mass range of 300–1750 m/z at a resolution of 70,000 FWHM and a target value of 3×10^6^ ions. Precursors were selected with an isolation window of 1.3 m/z (Q Exactive Plus) or 2.0 m/z (Q Exactive). HCD fragmentation was performed at a normalized collision energy of 25. MS/MS spectra were acquired with a target value of 10^5^ ions at a resolution of 17,500 FWHM, a maximum injection time (max.) of 55 ms and a fixed first mass of m/z 100. Peptides with a charge of +1, greater than 6, or with unassigned charge state were excluded from fragmentation for MS^2^, dynamic exclusion for 30s prevented repeated selection of precursors. The mass spectrometry proteomics data are available at the ProteomeXchange Consortium via the PRIDE partner repository with the data set identifier PXD015839.

#### Protein identification and quantification

Raw data were processed using MaxQuant software (version 1.5.7.4, http://www.maxquant.org/) (*42*) to calculate label-free quantification (LFQ) and iBAQ values (*43*). MS/MS spectra were searched for by the Andromeda search engine against a combined database containing the amino acid sequences of *Pto* (The Pseudomonas Genome Database) and 248 common contaminant proteins and decoy sequences. Trypsin specificity was required and a maximum of two missed cleavages were accepted. Minimal peptide length was set to seven amino acids. Carbamidomethylation of cysteine residues was set as fixed, oxidation of methionine and protein N-terminal acetylation as variable modifications. Peptide-spectrum matches and proteins were retained if they were below a false discovery rate of 1%.

#### Data analysis

Normalized iBAQ values were used for statistical analysis because the samples analyzed in the present study, which contain bacterial and plant proteins in different ratios, do not fulfill an assumption used in LFQ analysis that the abundance of the majority of proteins does not change between conditions. For simplification and comparison to transcriptome data, information on protein groups was omitted, and only the identifier displayed in the MaxQuant column “Fasta header” was used. For each sample, iBAQ values were normalized by TMM normalization in the package edgeR (*44*). For this, normalization factors were calculated with the calcNormFactors function with default settings based on proteins with iBAQ values greater than zero in all the samples. When iBAQ values were zero in more than one replicate out of three replicates, the protein was defined as “not detected” and the iBAQ values of all the replicates were converted to NA. TMM-normalized iBAQ values were then log_2_-transformed. To each protein, a linear model was fitted by using the function lmFit in the limma package with the following terms: Sgptr = GPTgpt + Rr+ ɛgtr, where S is the log_2_ count per million, GPT is the host genotype (or liquid medium): *Pto* strain: time point interaction and the random factors, R is the biological replicate, and ɛ is the residual. The eBayes function in the limma package was used for variance shrinkage in the calculation of the *p*-values, which was then used to calculate the false discovery rate (FDR; the Storey’s *q*-values) using the qvalue function in the qvalue package (*37*). To determine proteins with significant expression changes, the cutoff of *q*-value < 0.01 and |log_2_FC| >2 was applied. Hierarchical clustering was done using the dist and hclust functions in the R environment or using the Cluster3.0 software (*38*). Heatmaps were created with the TreeView (*39*). Enriched gene ontology terms were identified using BiNGO plugin for cytoscape (*40*).

#### Gene ontology (GO) analysis

mRNA/protein expression data were standardized using z-score and a GO expression matrix was generated by taking the mean z-score for each GO term. To select GO terms that show distinct expression patterns among different conditions, we performed statistical tests in all pairwise comparisons among 15 conditions for each GO term and manually curated GO terms with high numbers of significant pairs (redundant GO terms were avoided).

#### RT-qPCR analysis

RT-qPCR was performed using the SuperScript One-Step RT-PCR system kit (Invitrogen). As inputs, 30 ng of DNase-treated RNA extracted from infected leaves were used for analyzing bacterial genes.

#### Total protein extraction and immunoblotting analysis

*Pto* wild type and *Pto* AvrRpt2 strains were infiltrated into 4-week-old *A. thaliana* leaves and harvested at indicated time points. The infiltrated leaves were ground into fine powder in liquid nitrogen. Total proteins were extracted using the phenol extraction method. Briefly, equal amounts of protein extraction buffer (50 mM Tris-Cl pH 7.6, 5 mM EDTA, 5 mM EGTA, 2 mM DTT, and protease inhibition cocktail) and water-saturated phenol were added to ground tissues, and mixed well. The phenolic layer was moved to a new tube after centrifugation at 6,000 g for 5 min. Total proteins were precipitated in −20°C after adding an additional 2.5 volumes of ice-cold methanol containing 0.1 M ammonium acetate. Precipitated proteins were then twice washed in ice-cold methanol containing 0.1 M ammonium acetate and ice-cold 80% v/v acetone, respectively. Protein pellets were dissolved in protein sample buffer (40% v/v glycerol, 250 mM Tris-Cl pH 6.8, 3.5 M SDS, 5% v/v β-mercaptoethanol, 0.04% w/v bromophenol blue) after heating at 95°C for 5 min, and cooling down on ice for 1 min. For immunoblotting, proteins were separated on SDS-PAGE, and then transferred onto a polyvinylidene difluoride membrane using an electrophoretic apparatus. Protein detection was performed using anti-HrcC (1:5000) (*45*) and anti-HrpZ (1:5000) (*46*) as primary antibodies, and anti-mouse (1:10000) and anti-rabbit (1:10000) conjugated with horseradish peroxidase as secondary antibodies. To compare the expression of HrpZ between samples, protein loading was first normalized to HrcC expression.

## Supplementary Text

### Conditions used in this study

Note that this is not a time-course study as the doses of starting bacterial inocula are different for the two time points (OD_600_=0.5 for 6 hpi and OD_600_=0.005 for 48 hpi). *Pto* AvrRpt2 was not used for the sampling at 48 hpi because this strain caused tissue collapse in the leaves in this condition.

### Bacterial proteins differentially expressed under various conditions (related to Fig. S1)

We analyzed mRNAs and proteins whose expression was significantly changed between the *in vitro* (KB) and *in planta* (Col-0) conditions at 6 hpi or 48 hpi. GO enrichment analysis showed that mRNAs as well as proteins related to “pathogenesis”, “translation”, and “cell wall organization or biogenesis” were induced both at 6 and 48 hpi (Cluster I and II in Fig. S1A and Cluster I in Fig. S1B). Interestingly, in the proteome data, tRNA synthases and ribosomal proteins, both of which are related to protein translation, showed the opposite expression patterns, i.e., tRNA synthases were suppressed at 48 hpi, while ribosomal proteins were induced (Cluster I and III in Fig. S1B). Whether bacterial translation efficiency is reduced or enhanced at 48 hpi compared with the other conditions remains to be elucidated. mRNAs and proteins annotated as “response to oxidative stress” were induced at 48 hpi (Cluster III in Fig. S1A and cluster IV in Fig. S1B), implying that *Pto* experienced oxidative stress at a later stage of infection or that a longer exposure to oxidative stress caused such induction. A substantial number of *Pto* mRNAs (324 mRNAs) and proteins (196 proteins) were differentially expressed among different host genotypes at 48 hpi (Fig. S1C and S1D), whereas host genotype effects were small at 6 hpi (only 15 mRNAs and one protein were differentially expressed, respectively). *Pto* AvrRpt2, which was strongly affected by host SA pathways at 6 hpi at the transcriptome level (*8*), showed similar proteome patterns among SA mutant plants (only one protein was differentially expressed). This might be because transcriptional changes were not yet reflected in protein accumulation at 6 hpi. Thus, we focused on analyzing host genotype effects on protein expression in *Pto* at 48 hpi. There were 196 proteins whose expression was significantly affected in at least one of the SA mutants (Fig. S1D). Gene ontology (GO) enrichment analysis revealed that pathogenesis-related proteins (“Interaction with host”) were highly expressed in the SA mutants (Cluster II in Fig. S1D), suggesting that the SA immune pathway suppresses the expression of pathogenesis-related proteins. On the other hand, expression of “translation”-related proteins (ribosomal proteins) was suppressed in the SA mutants (Cluster I in Fig. S1D). Taken together, the SA pathway affects expression of bacterial proteins related to bacterial virulence and basic metabolism *in planta*.

### GO analysis

As shown in Fig. S1, bacterial functions differentially expressed in different conditions could be studied by GO enrichment analysis following statistical tests. However, this analysis is highly dependent on the thresholds applied for selecting differentially expressed mRNAs or proteins, and thus some important information can be lost before GO enrichment analysis. Moreover, it is difficult to compare the global expression pattern of GO terms across many conditions in this approach. To gain insights into biological functions that are differentially regulated in various conditions, we annotated the *Pto* genome with GO terms and calculated GO expression scores (see Materials and Methods). This enabled quantitative analysis of bacterial functions across various conditions (Fig. 2).

## Supplementary figures

**Fig. S1:**
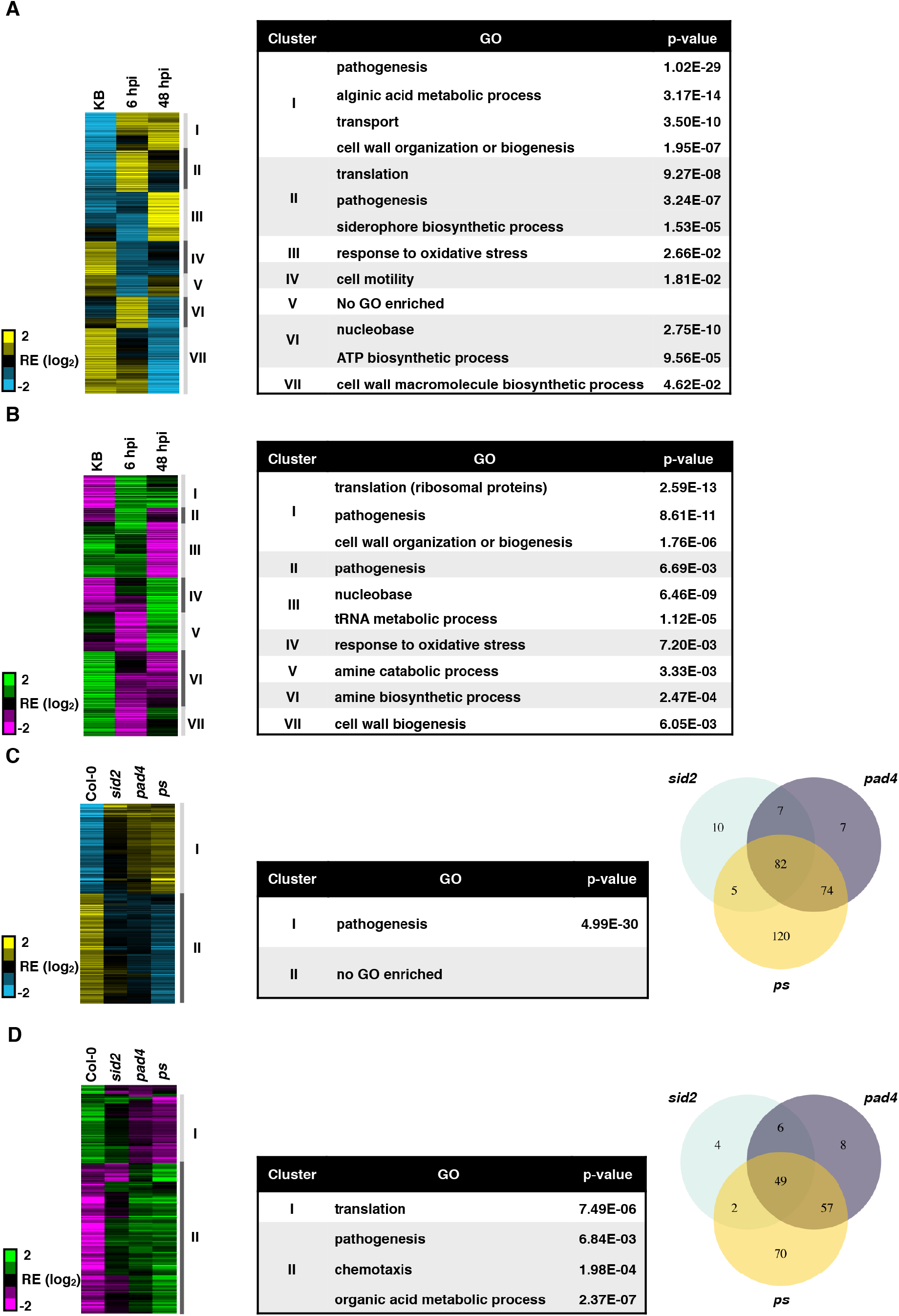
Distinct patterns of *Pto* transcriptomes and proteomes in various conditions. (A and B) Hierarchical clustering of the relative expression (RE) of *Pto* mRNAs (A) and proteins (B) differentially expressed in at least one condition among *in vitro* (King’s B medium (KB)) and *in planta* conditions (Col-0) at 6 hpi and 48 hpi (FDR < 0.01; |log_2_FC| > 1). List of gene ontology (GO) terms enriched in the clusters shown in the sidebar of the heatmap. (C and D) Hierarchical clustering of the relative expression of *Pto* mRNAs (C) and proteins (D) differentially expressed in at least one plant SA mutant compared with the wild-type Col-0 at 48 hpi (FDR < 0.01; |log_2_FC| > 1). List of GO terms enriched in the clusters shown in the sidebar of the heatmap. Venn diagrams of *Pto* mRNAs/proteins differentially expressed in each plant mutant compared with Col-0. For the mRNA/protein expression data and full GO lists, see Data S3-S7. *ps*, *pad4 sid2*.

**Fig. S2:**
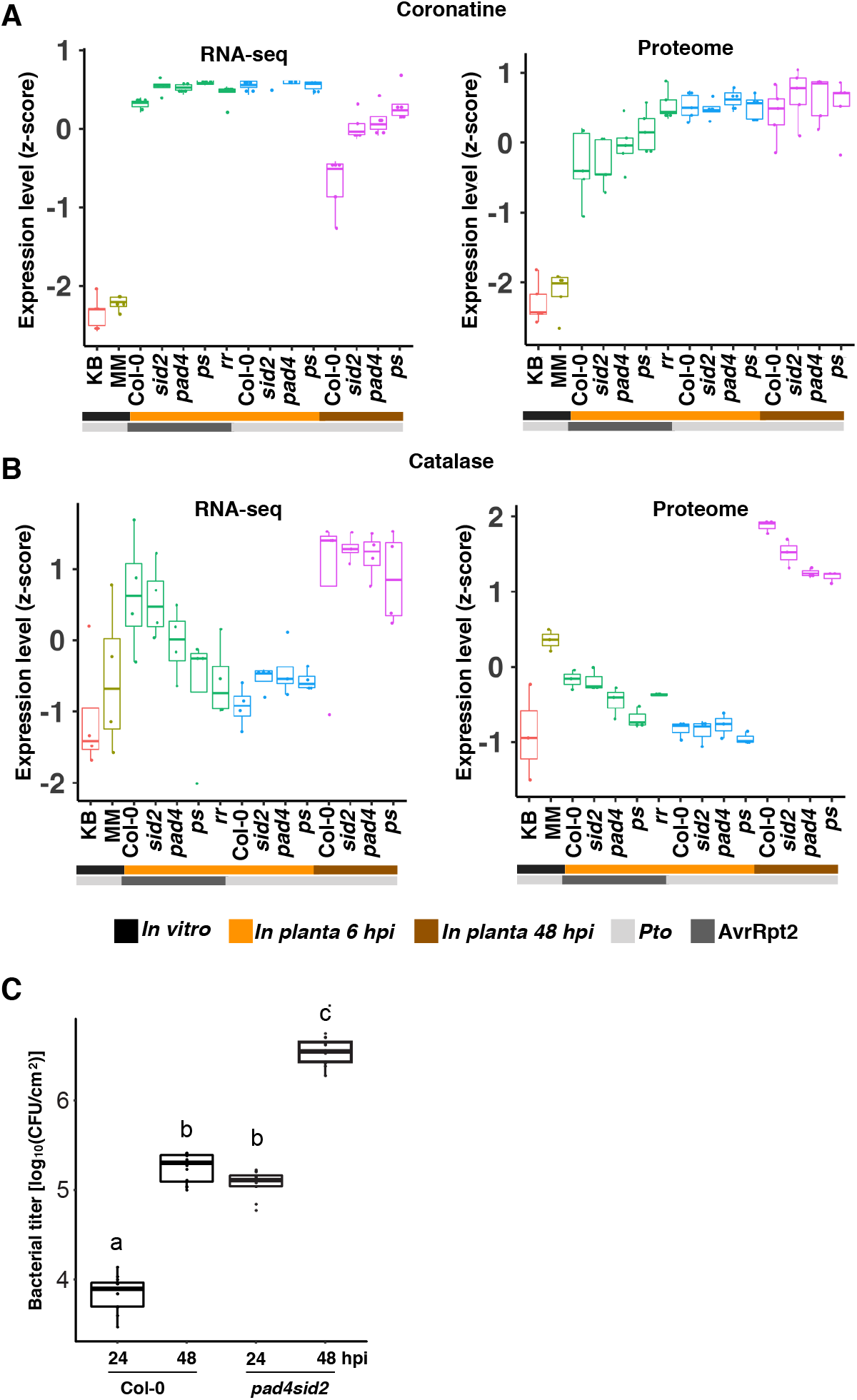
Regulation of selected mRNAs and proteins. Box plots show the expression (z-score) of mRNAs (left) and proteins (right) related to “coronatine biosynthesis” and “catalase” under various conditions. Light and dark gray sidebars represent *Pto* and *Pto* AvrRpt2, respectively. Black, orange, and brown sidebars represent *in vitro* (KB), *in planta* (Col-0) 6 hpi, and *in planta* 48 hpi, respectively. MM, minimal medium; *ps*, *pad4 sid2*; *rr*, *rpm1 rps2*.

**Fig. S3:**
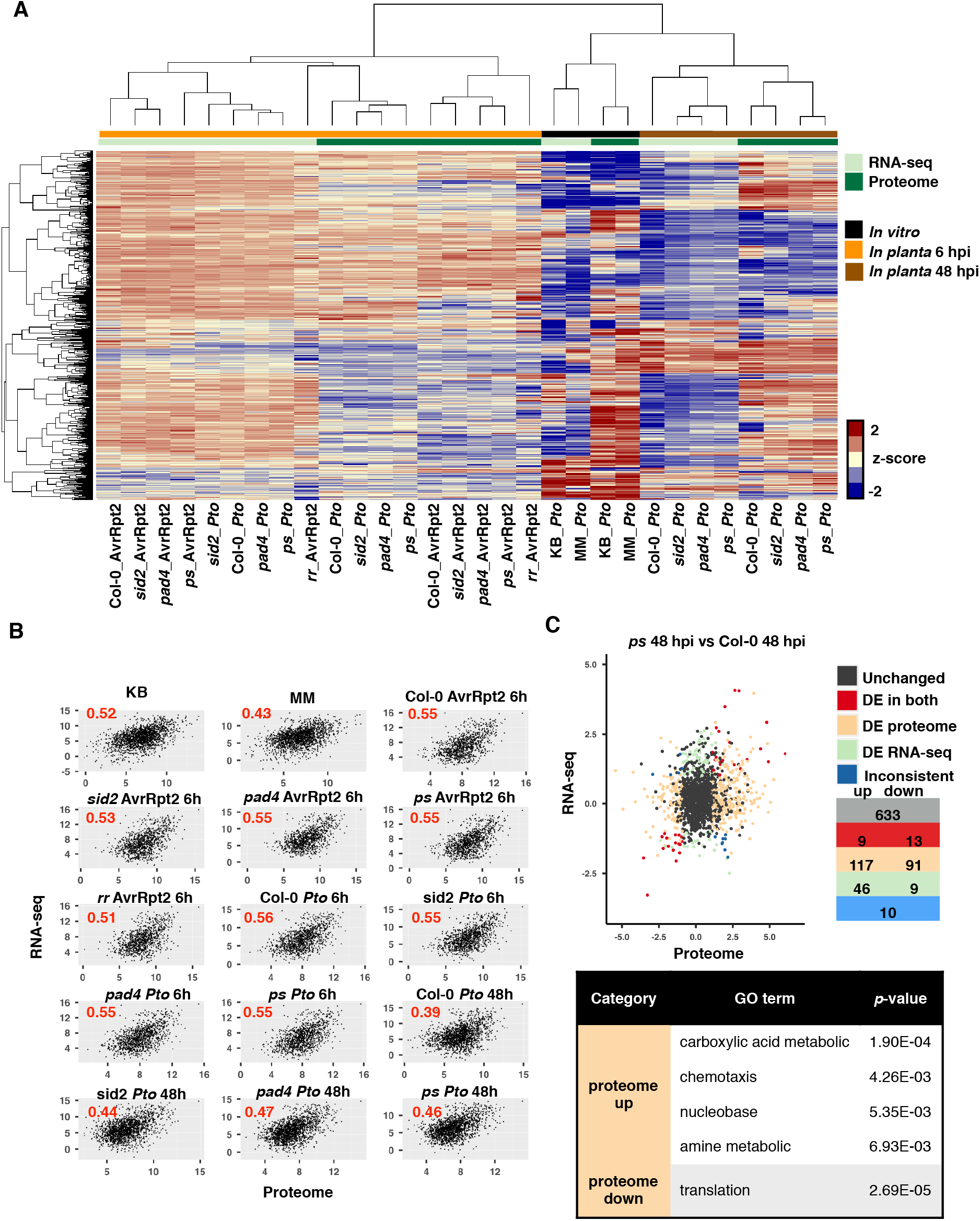
Integration of bacterial transcriptome and proteome data. (A) Transcriptome and proteome data were standardized using z-scores (log_2_) and combined, followed by hierarchical clustering. mRNAs/proteins detected in all conditions were subject to analysis. Light and dark green sidebars represent transcriptome and proteome data, respectively. Black, orange, and brown sidebars represent *in vitro* (King’s B (KB)), *in planta* (Col-0) 6 hpi, and *in planta* 48 hpi, respectively. MM, minimal medium; *ps*, *pad4 sid2*; *rr*, *rpm1 rps2*. (B) Comparisons between transcriptome and proteome data in each condition. Pearson’s correlation coefficients were shown. (C) mRNA/protein expression fold changes between Col-0 and *pad4 sid2* at 48 hpi were compared. mRNAs/proteins differentially expressed (DE; FDR < 0.01; |log_2_FC| > 2) in both, either, and neither (“Unchanged”) of the transcriptome and proteome studies were grouped and colored. mRNAs/proteins differentially expressed in the opposite direction were colored in blue (“Inconsistent”). The numbers of mRNAs/proteins are shown for each category. List of gene ontology (GO) terms enriched in the group of proteins that are significantly induced *in planta* at both mRNA and protein levels or proteins that are significantly suppressed *in planta* only at the protein level. For the full gene list and GO list, see Data S10 and S11. (B and C) mRNAs/proteins detected in both the transcriptome and proteome in each condition or comparison were used for this analysis.

**Fig. S4:**
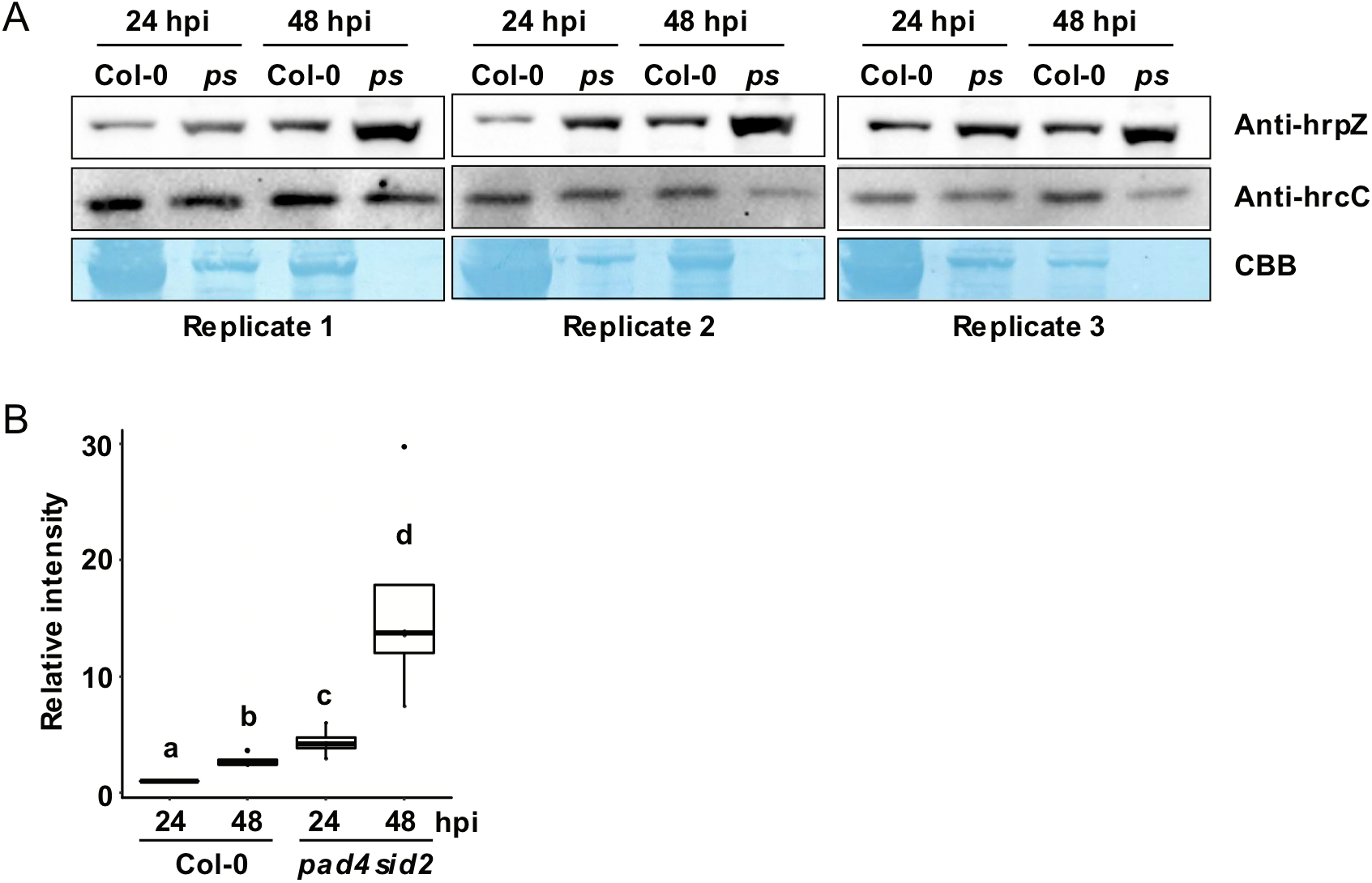
Component-specific targeting of the type III secretion system by plant SA pathways. (A) HrpZ and HrcC proteins were detected by immunoblotting using specific antibodies in Col-0 and *pad4 sid2* (*ps*) at 24 hpi and 48 hpi. Protein loading amount was adjusted to the expression of the HrcC protein. (B) Relative intensity of HrpZ expression normalized to HrcC expression based on the data in (A). Different letters indicate statistically significant differences (adjusted P < 0.05; Benjamini–Hochberg method)

**Fig. S5:**
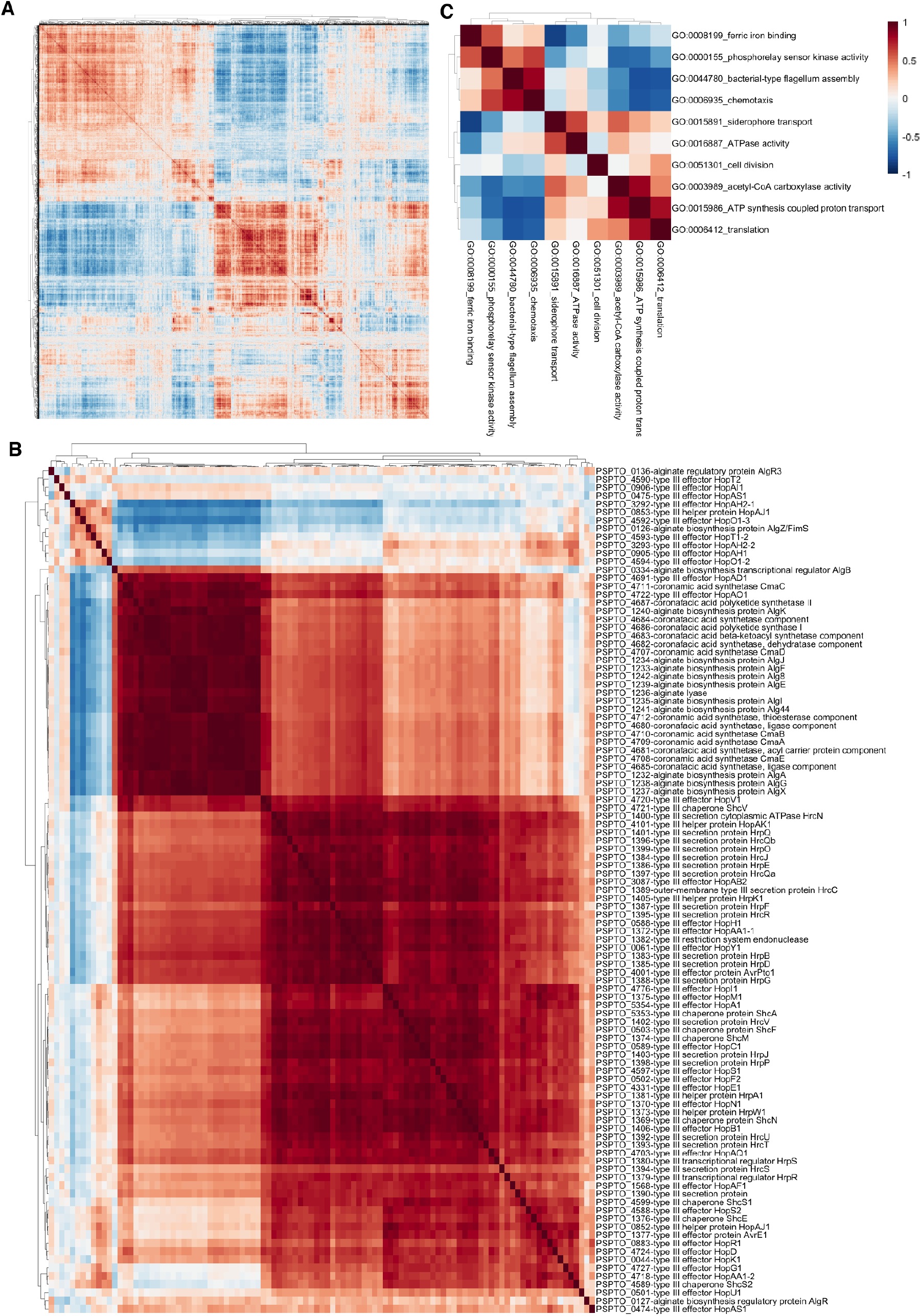
Co-expression analysis of *Pto* mRNAs. (A) Correlation matrix of 4,765 *Pto* mRNAs based on 125 transcriptome datasets in 38 conditions. (B) Correlation matrix of expression of mRNAs related to coronatine, alginate, and the type III secretion system (T3SS) under various conditions. (C) Correlation matrix of bacterial gene ontology (GO) expression. Selected GOs are shown.

**Fig. S6:**
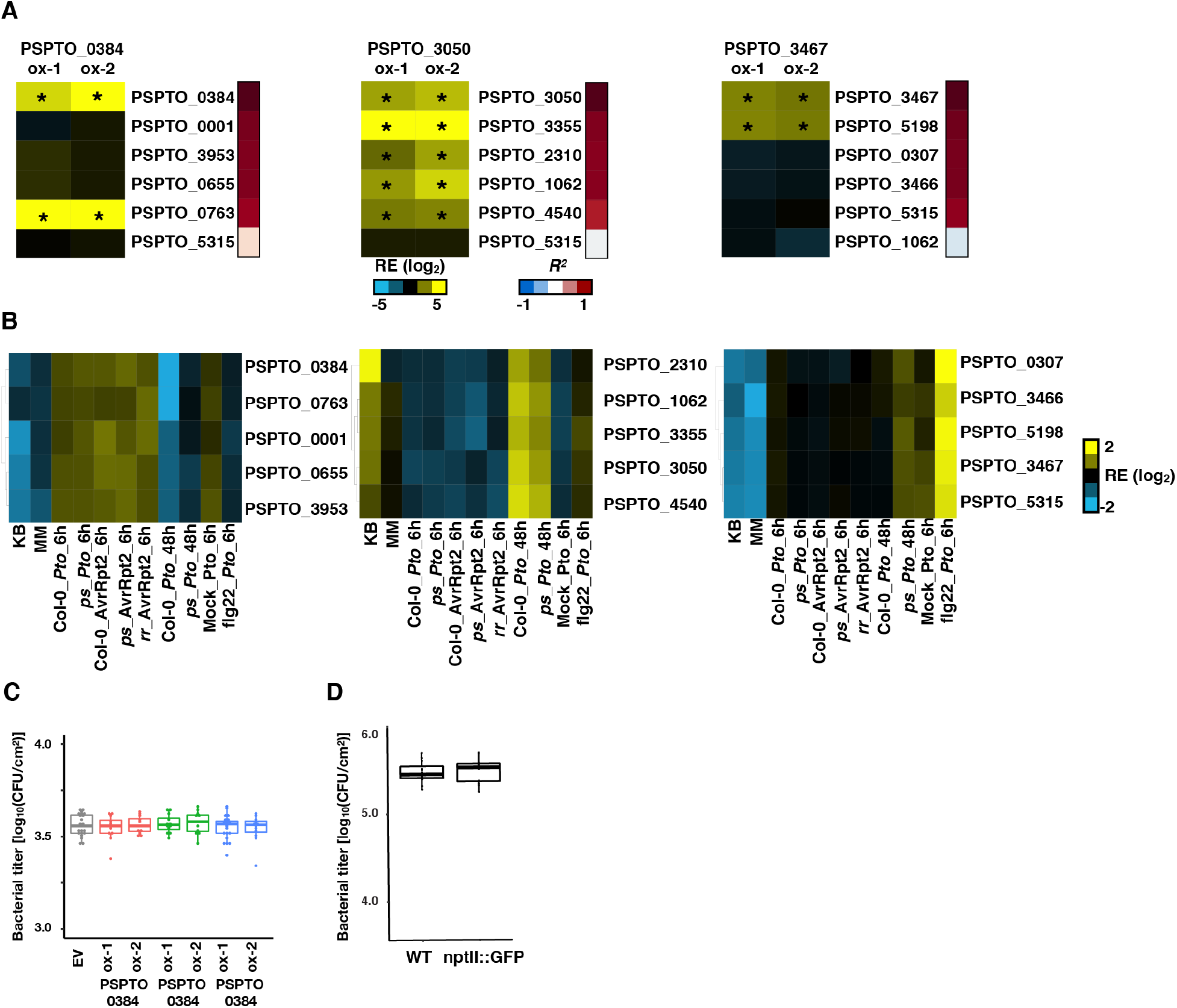
mRNA co-expression analysis of *Pto* reveals functional mRNA clusters and their regulatory logic. (A) *In planta* (Col-0) mRNA expression fold changes between *Pto* overexpressing putative transcriptional regulators (TRs) and wild type *Pto*. Asterisks indicate significant differences (|log_2_FC| >2, *p*-value < 0.05). Correlation coefficients between each mRNA and the TRs are shown in the sidebars on the right. (B) Relative expression (RE) of the TRs and mRNAs highly co-expressed with each of the TRs. (C) Growth of *Pto* overexpressing putative TRs or GFP (infiltrated at OD_600_ = 0.001) in Col-0 at 0 hpi. n = 12 biological replicates from three independent experiments. (D) Growth of wild type *Pto* and *Pto* overexpressing GFP (infiltrated at OD_600_ = 0.001) in Col-0 at 48 hpi. n = 12 biological replicates from three independent experiments. Overexpression of GFP did not affect the growth of *Pto in planta*.

## Supplementary Data

**Data S1.** Full RNA-seq dataset.

**Data S2.** Full proteome dataset.

**Data S3.** RNA-seq or proteome data shown in Fig. S1A–D.

**Data S4.** Full GO list for Fig. S1A.

**Data S5.** Full GO list for Fig. S1B.

**Data S6.** Full GO list for Fig. S1C.

**Data S7.** Full GO list for Fig. S1D.

**Data S8.** GO-based RNA-seq data shown in Fig. 2A.

**Data S9.** GO-based proteome data shown in Fig. 2B.

**Data S10.** Full GO list for Fig. 3B.

**Data S11.** Full GO list for Fig. 3C.

**Data S12.** Gene expression data used for co-expression analysis in Fig. 4A

**Data S13.** Correlation matrix shown in Fig. 4A.

**Data S14.** Primers used in this study.

